# Transcription-induced formation of extrachromosomal DNA during yeast ageing

**DOI:** 10.1101/744896

**Authors:** Ryan Hull, Michelle King, Grazia Pizza, Felix Krueger, Xabier Vergara, Jonathan Houseley

## Abstract

Extrachromosomal circular DNA (eccDNA) facilitates adaptive evolution by allowing rapid and extensive gene copy number variation, and is implicated in the pathology of cancer and ageing. Here, we demonstrate that yeast aged under environmental copper accumulate high levels of eccDNA containing the copper resistance gene *CUP1*. Transcription of *CUP1* causes *CUP1* eccDNA accumulation, which occurs in the absence of phenotypic selection. We have developed a sensitive and quantitative eccDNA sequencing pipeline that reveals *CUP1* eccDNA accumulation on copper exposure to be exquisitely site specific, with no other detectable changes across the eccDNA complement. eccDNA forms *de novo* from the *CUP1* locus through processing of DNA double-strand breaks (DSBs) by Sae2 / Mre11 and Mus81, and genome-wide analyses show that other protein coding eccDNA species in aged yeast share a similar biogenesis pathway. Although abundant we find that *CUP1* eccDNA does not replicate efficiently, and high copy numbers in aged cells arise through frequent formation events combined with asymmetric DNA segregation. The transcriptional stimulation of *CUP1* eccDNA formation shows that age-linked genetic change varies with transcription pattern, resulting in gene copy number profiles tailored by environment.

## Introduction

In contrast to the normally sedate evolution of chromosomal DNA, eccDNA can be rapidly accumulated and lost in eukaryotic cells, facilitating timely changes in gene expression and accelerating adaptation. eccDNA accumulation provides a pathway for adapting to drug treatment, environmental stress and genetic deficiency in diverse eukaryotes [1–5]. Amplification of both driving oncogenes and chemotherapy resistance genes on eccDNA has also been widely reported in tumour cells and is a frequent feature of cancer genomes [6, 7]. Genome-wide studies in yeast, worms and non-transformed mammalian cells reveal huge diversity in the protein coding eccDNA complement [8–10], in addition to widespread microDNA, heterochromatic repeat-derived eccDNA, and telomeric circles [11–13]. Nevertheless, we understand little of the mechanisms of protein-coding eccDNA formation as such events are generally rare and unpredictable.

Re-integration of eccDNA provides an efficient pathway for gene amplification, with various chromosomal adaptations in lower eukaryotes being known or predicted to have emerged through eccDNA intermediates [14–18]. In tumour cells, aggregation of smaller eccDNA to large (1-5MB), microscopically visible double minutes, and re-integration in homogeneously staining regions to yield high-copy chromosomal repeats has also been reported [7, 19, 20]. Thereby, eccDNA acts as an intermediate in chromosome structural evolution. However, eccDNA usually segregates randomly during mitosis due to the absence of a centromere, causing pronounced heterogeneity in eccDNA complement across the population that provides dramatic phenotypic plasticity [21]. Theoretical models predict that gene copy number amplification can occur much faster on eccDNA than chromosomal DNA [7], and population heterogeneity in gene dosage on eccDNA can be selected not only for increased gene dosage [21, 22], but also for decreased dosage due to the ease of eccDNA loss in cell division [18, 23, 24].

Segregation of eccDNA during cell division is not necessarily random [25]. This is exemplified by budding yeast replicative ageing wherein cells divide asymmetrically into mother and daughter cells, with mothers retaining various molecules that could be detrimental to daughter cell fitness, including a well-studied eccDNA species - extrachromosomal ribosomal DNA circles (ERC) [26, 27]. Retention of ERCs in mother cells involves tethering by SAGA and TREX2 to nuclear pore complexes, which are themselves largely retained in mother cells, with mutation of SAGA components abrogating retention and extending mother cell lifespan [28–30]. ERCs replicate on each division and mother cell retention leads to exponential copy number amplification, increasing genome size by 50% in 24 hours [31], which is thought to interfere with various critical pathways to accelerate, if not cause, ageing pathology [32, 33].

Formation mechanisms for eccDNA have been primarily inferred from amplicon structure. The conceptually simple episome model invokes a recombination event between distal sites on the same chromosome to yield an eccDNA with a matching deletion [19]. Chromosomal deletions closely matching eccDNA breakpoints have been observed although not universally [3, 19, 34–37]. However, recombination would be expected to favour sites with substantial homologous sequence, and although this is observed at breakpoints in some systems, microhomology is commonly reported [3, 9, 12, 22, 37, 38]. Conversely, replication-based models allow eccDNA formation without chromosomal change. Substantial evidence supports the formation of ERCs in yeast from replication forks stalled at the ribosomal DNA (rDNA) replication fork barrier [39, 40], however it is unclear if such events are specific to the yeast rDNA which contains a unique replication fork stalling system [41, 42]. Other mechanisms invoke re-replication of DNA to yield nested structures that must be resolved by homologous recombination prior to mitosis [43], and can potentially yield eccDNA of great complexity. Such models predict no clear structural signature and have therefore been hard to test, although derepression of origin re-licensing does lead to chromosomal copy number variation as predicted [44]. Without reproducible methods to observe eccDNA formation *de novo*, it remains very difficult to validate these models or probe mechanistic details.

A few examples of regulated eccDNA formation have been reported. Stage specific *de novo* formation of eccDNA during embryonic development in Xenopus facilitates rolling circle amplification of the rDNA [45, 46], while eccDNA production in T- and B-cells as a side-effect of highly-regulated V(D)J recombination is well-studied [47]. In yeast, rDNA recombination including ERC formation is highly dependent on histone deacetylases that respond to nutrient availability [4, 48, 49], leading us to question whether protein-coding eccDNA formation could also be driven by environmentally-responsive signalling, analogous to the stimulated chromosomal copy number variation that we recently reported [50]. Here we describe the transcriptionally-stimulated formation of eccDNA from the yeast *CUP1* locus and characterise the mechanism by which eccDNA is processed from DSBs.

## Results

### Transcription promotes eccDNA accumulation from the CUP1 locus

ERCs are enriched in aged yeast and we speculated that other eccDNA may be similarly enriched during ageing. To obtain highly purified aged yeast samples we employed the mother enrichment program (MEP) combined with cell wall labelling [51, 52]. Young cells in log phase growth are labelled with a cell wall reactive biotin derivative that is completely retained on mother cells since daughter cell walls are newly synthesised on each division [52]. Estradiol is added to activate the MEP, after which all new-born daughter cells become non-replicative. Mother cells are therefore the only replicative cells in the population, and reach advanced age without the culture becoming saturated; these highly-aged mother cells can then be affinity purified (Figure 1A).

**Figure 1:**
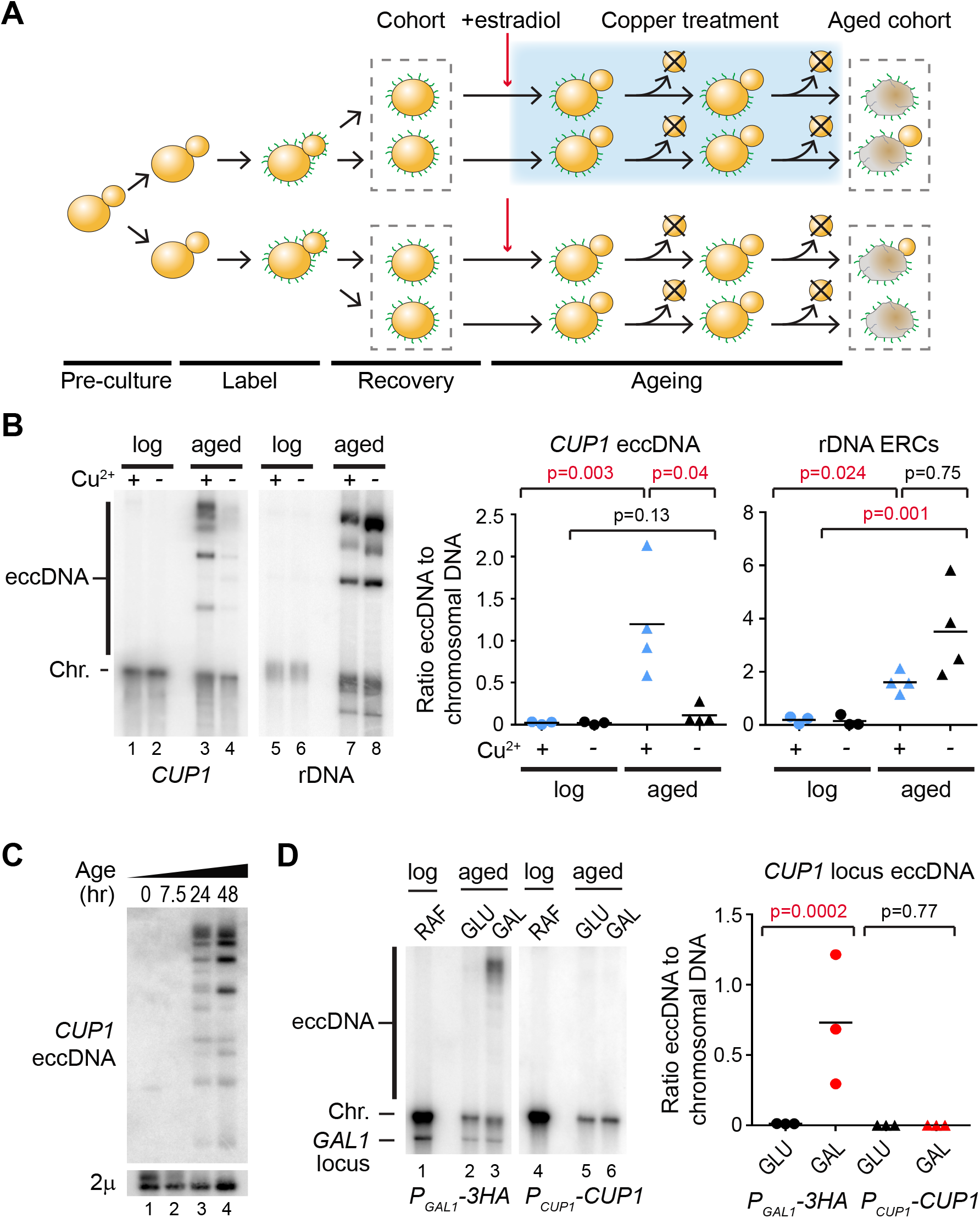
Transcription of the *CUP1* locus causes eccDNA accumulation. A: Schematic representation of cell labelling, induction and ageing in the presence or absence of copper. B: Southern blot analysis of *CUP1* eccDNA and ribosomal DNA-derived eccDNA (ERCs) in yeast cells aged for 48 hours in the presence or absence of 1mM CuSO_4_, along with young cells maintained in log phase in the presence or absence of 1mM CuSO_4_ for an equivalent time. Large linear fragments of chromosomal DNA (Chr.) migrate at the resolution limit of the gel, while circular DNA species (eccDNA) migrate more slowly. Abundances of eccDNA and ERCs were compared by one-way ANOVA, n=4 biological replicates, data was log-transformed for testing to fulfil the assumptions of a parametric test. C: Southern blot analysis of *CUP1* eccDNA in cells aged for 0, 7.5, 24 or 48 hours in 1mM CuSO_4_. Chromosomal DNA was removed with ExoV to improve sensitivity, endogenous circular 2μ DNA is shown as a loading control. D: Southern blot analysis of eccDNA in heterozygous strain bearing one *P_GAL1_-3HA cup1* locus modified by replacing all *CUP1* promoters and ORFs with *P_GAL1_* promoters and 3HA ORFs, and one wild-type *P_CUP1_-CUP1* allele. eccDNA is detected with allele specific probes, additional band in the left panel is from hybridisation to the endogenous *GAL1* locus on chromosome II. Quantification and analysis performed as in B, n=3.

To test the potential for environmentally-stimulated eccDNA accumulation, we analysed the locus encoding the copper-resistance gene *CUP1*. Exposure to environmental copper strongly induces transcription of the *CUP1* gene, which is encoded in an array of 2-15 2kb tandem repeats on chromosome VIII [53, 54]. A labelled cohort of cells was aged in the presence or absence of 1mM CuSO_4_ for 48 hours (Figure 1A lanes 3,4), and control cells were maintained at log phase for the same period. Southern blot analysis of aged cohorts revealed multiple DNA species in the copper-treated samples migrating above the resolution limit of the gel, a region that is largely devoid of linear DNA (Figure 1B lane 3). These were much less prominent for cells aged in the absence of copper, and undetectable in log phase cells (Figure 1B lanes 1-4).

To confirm that these signals represent eccDNA, genomic DNA from cells aged in 1mM CuSO_4_ was digested with exonuclease V (ExoV), which degrades linear DNA progressively from double stranded ends, but has no activity on circular or nicked circular DNA. As expected, ExoV efficiently degraded DNA from the chromosomal *CUP1* band while the species above the resolution limit were unaffected (Figure S1). Copper-stimulated formation of *CUP1* eccDNA was reproducible and significant, and did not represent a general change in eccDNA accumulation as ERCs were detected at a slightly, but not significantly, higher level in cells aged in the absence of copper (Figure 1B, lanes 5-8). A timecourse analysis was then performed using genomic DNA treated with ExoV, which increases sensitivity by removing chromosomal DNA signals, revealing that *CUP1* eccDNA is readily detected after 24 hours in copper, but not after 7.5 hours, so eccDNA accumulation increases with age (Figure 1C). Although eccDNA signals are stronger after 48 hours, we have previously noted that Cu^2+^ reduces cell viability during ageing [50], and we therefore restricted further experiments involving ageing cells in CuSO_4_ to 24 hours.

Accumulation of *CUP1* eccDNA may offer a selective advantage by increasing copper resistance. To ensure that selection for copper resistance does not underlie the observed eccDNA accumulation, we performed a similar experiment using a strain in which transcription at the *cup1* locus is induced under a different environment and does not yield active protein. We have previously developed a yeast strain in which all *CUP1* repeats have been mutated such that the *CUP1* protein-coding sequences are replaced with a non-functional *3HA* sequence and the *P_CUP1_* promoters are replaced by galactose-responsive *GAL1* promoters (*P_GAL1_*) [50]. These cells transcribe mRNA encoding only a non-functional 3HA protein from every *cup1* repeat in the presence of galactose but not glucose. Here, we mated this strain with a wild-type haploid to form a heterozygous diploid with one galactose-responsive *P_GAL1_-3HA cup1* allele and one copper responsive *P_CUP1_-CUP1* allele. When aged in galactose, we observed that these cells form eccDNA from the *P_GAL1_-3HA* allele but not from the copper-responsive wild-type *P_CUP1_-CUP1* allele, whereas no eccDNA from either allele was detectable in cells aged in glucose (Figure 1D, compare lanes 2,3 and 5,6).

These experiments show that transcriptional activiation of the *CUP1* locus in aged cells causes the accumulation of eccDNA even in the absence of selection, and that eccDNA accumulation is specific to the transcriptionally induced locus versus a non-induced control locus.

### eccDNA accumulation is locus specific

To reveal eccDNA distribution across the genome, we developed a quantitative eccDNA sequencing protocol for the ~30ng genomic DNA available from our standard ageing yeast preparations. Previous eccDNA sequencing analysis in yeast has involved exonuclease digestion and rolling circle amplification [9]. The latter reaction is affected by circle size, making quantitative comparisons between circular species challenging. Alternative biochemical methods are quantitative but require a prohibitive amount of starting material [10]. We therefore employed extensive exonuclease treatment of aged genomic DNA followed by DNA fragmentation and a sensitive library preparation protocol to reveal the circular DNA complement (Figure S2A).

Exonuclease treatment alone enriched for ERCs and sub-telomeric eccDNA (Figure S2A), however >95% of sequencing reads in these libraries mapped to rDNA and *2*μ, which limited sensitivity. To remove these species, each sample was split in three parts and treated with different restriction enzymes prior to exonuclease treatment. All three restriction enzymes cleave ERCs and 2μ, rendering both sensitive to exonuclease digestion, whereas any circle that does not contain all three restriction sites will be maintained in at least one reaction (Figure S2B). After exonuclease treatment, the three parts were reunited for sequencing. This Restriction-digested Extrachromosomal Circular DNA sequencing protocol (REC-seq) (Figure 2A) reduced reads from ERCs and 2μ to <2%, revealing eccDNA from other regions including Ty elements, *CUP1*, the *ENA* locus and *HXT6-HXT7*, all of which have been previously described using less quantitative methods (Figure S2B) [9].

**Figure 2:**
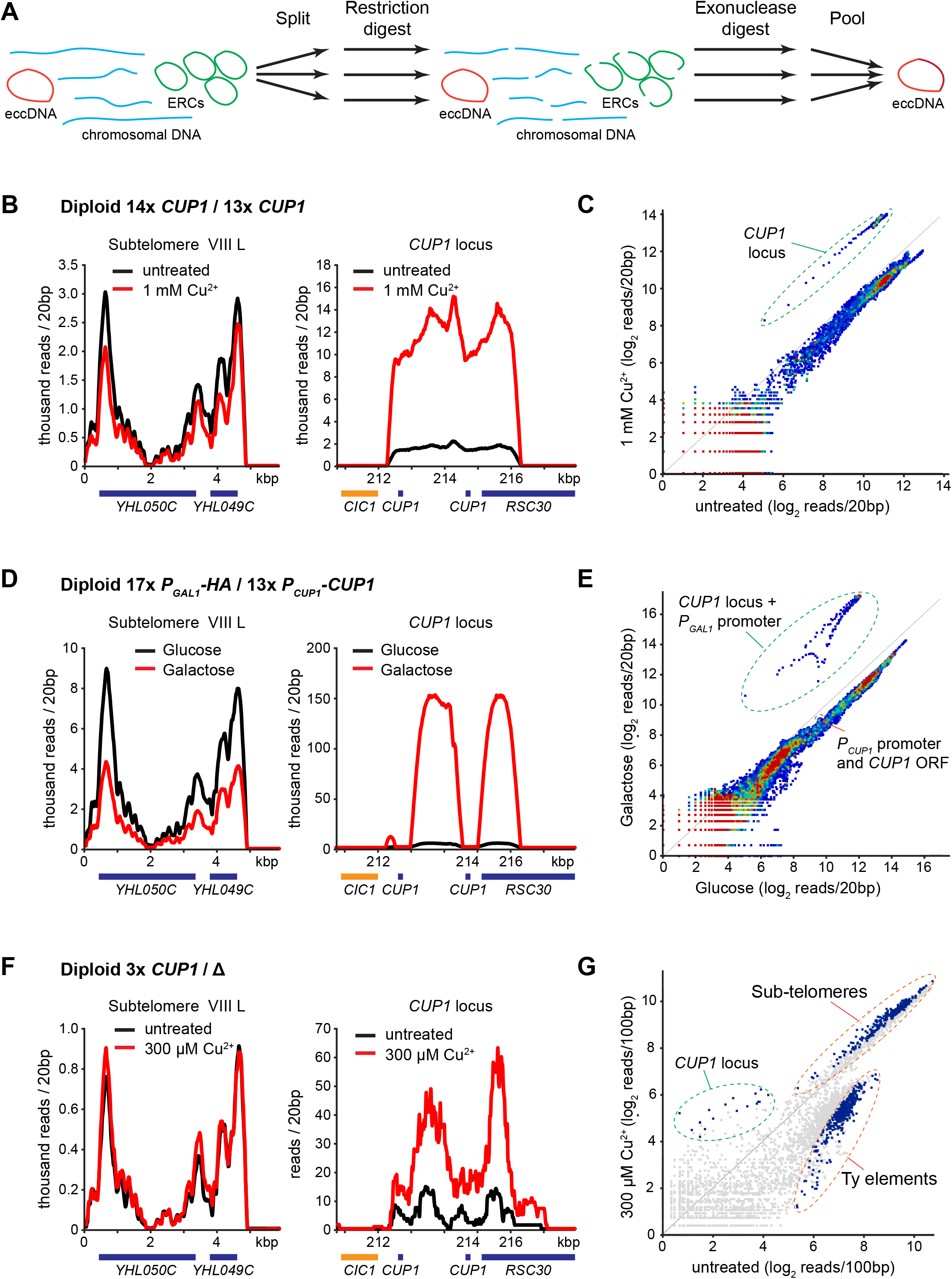
Genome-wide quantitative mapping of eccDNA in aged cells. A: Schematic representation of library preparation method, briefly genomic DNA is split into three samples and digested with three different endonucleases that all target ERCs and 2μ DNA, then digested with ExoV and ExoI. After two rounds of treatment, samples are re-united, fragmented and a DNA sequencing library prepared by standard methods. B: Distribution of sequencing reads around the *CUP1* locus on chromosome VIII and sub-telomere VIII L obtained from REC-seq of cells aged for 24 hours in the presence or absence of 1mM CuSO_4_. C: Scatter plot of read counts in 20 bp bins across the genome for samples shown in B. Circled points represent bins within the *CUP1* repeat region. D&E: Equivalent analysis to B&C performed on *P_GAL1_-3HA / P_CUP1_-CUP1* heterozygous strain aged for 48 hours in glucose or galactose. The pronounced dip in reads mapping to the *CUP1* locus over the actual *CUP1* ORFs corresponds to the region of the *CUP1* repeat replaced with *P_GAL1_-3HA*, and demonstrates that essentially all the eccDNA is formed from the *P_GAL_-HA* allele. Bins representing the *CUP1* promoter and ORF are also highlighted in E. F: Analysis of reads mapping to *CUP1* locus and sub-telomere VIII L averaged across 3 biological replicates of 3x*CUP1* / Δ cells aged for 24 hours in the presence or absence of 300μM CuSO_4_. G: Scatter plot of average read counts across 3 biological replicates in 100 bp bins across the genome for samples shown in F. Blue dots are significantly different between cells aged in the presence or absence of copper based on edgeR analysis, p<0.05. The genomic location of the different clusters of significantly different bins are indicated, all those within the *CUP1* locus area are derived from *CUP1* and are the most enriched in the presence of copper.

In agreement with Southern blotting data, REC-seq on cells aged in the presence of copper showed a substantial (~10-fold) increase in reads mapping to the *CUP1* locus but not to other loci such as the sub-telomeric regions (Figure 2B). Read quantification across the genome in 20bp windows revealed that although eccDNA accumulates from many loci in aged cells, only the *CUP1* locus is differentially affected after 24 hours (Figure 2C). Applying the same method to the heterozygous *P_GAL1_-3HA* / *P_CUP1_-CUP1* strain aged for 48 hours in galactose or glucose media revealed many more *CUP1* reads in cells aged on galactose, with a pronounced dip in the profile at the region of the repeat containing the *CUP1* promoter and ORF, both of which are absent from the *P_GAL1_-3HA* allele but present in the *P_CUP1_-CUP1* allele (Figure 2D). Again, by scatter plot the specificity of eccDNA accumulation from the mutated *P_GAL1_-3HA cup1* allele was obvious, while reads mapping to the *CUP1* promoter and ORF that are only present on the wild-type *P_CUP1_-CUP1* allele were not differentially enriched (Figure 2E). Therefore, eccDNA accumulation from the *CUP1* locus promoted by transcription is remarkably locus specific.

REC-seq is more sensitive than Southern blot, allowing quantification of *CUP1* eccDNA accumulation in a strain with only three tandem copies of the *CUP1* locus; this represents a more general case as loci containing small numbers of tandem repeats are common in eukaryotic genomes [55, 56]. eccDNA from *CUP1* accumulated more in cells aged in copper (Figure 2F); the difference (~4-fold) was smaller than in the MEP wild-type strain, but this is not unexpected given that the 3x*CUP1* / Δ cells contain 9-fold fewer copies of *CUP1*. The genome-wide profile of eccDNA was more affected by copper treatment in these cells, likely due to greater copper stress, and many eccDNAs accumulated to lower levels (Figure 2G). However, edgeR analysis of differential accumulation across 3 biological replicates revealed that the significant regions most over-represented in the copper-treated sample all derive from the *CUP1* locus. Additionally, Ty-element eccDNA was under-accumulated in copper treated cells, while sub-telomeric eccDNA was over-accumulated though by a very small amount.

Therefore, eccDNA accumulation caused by transcription is remarkably site specific and is observable even in low-copy tandem repeats.

### Asymmetric segregation is necessary but not sufficient for eccDNA accumulation

Asymmetric segregation of replicating ERCs causes ERC accumulation during ageing. The retention of ERCs in mother cells depends on attachment to the nuclear pore complex through the SAGA chromatin modifying complex, and rapid progression from S phase through mitosis so that ERCs are not passed to daughter cells by diffusion [28, 29]. We found that cells lacking the SAGA component Spt3 contain far less *CUP1* eccDNA and ERCs after 24 hours (Figure 3A), even though transcriptional induction of the *CUP1* promoter was normal in *spt3*Δ cells (Figure S3). We observed a similar defect in *bud6*Δ and *yku70*Δ mutants that are defective in ERC asymmetric segregation due to bud neck formation issues or extended G2 respectively [29, 57](Figure 3B). Therefore, asymmetric segregation is necessary for *CUP1* eccDNA accumulation.

**Figure 3:**
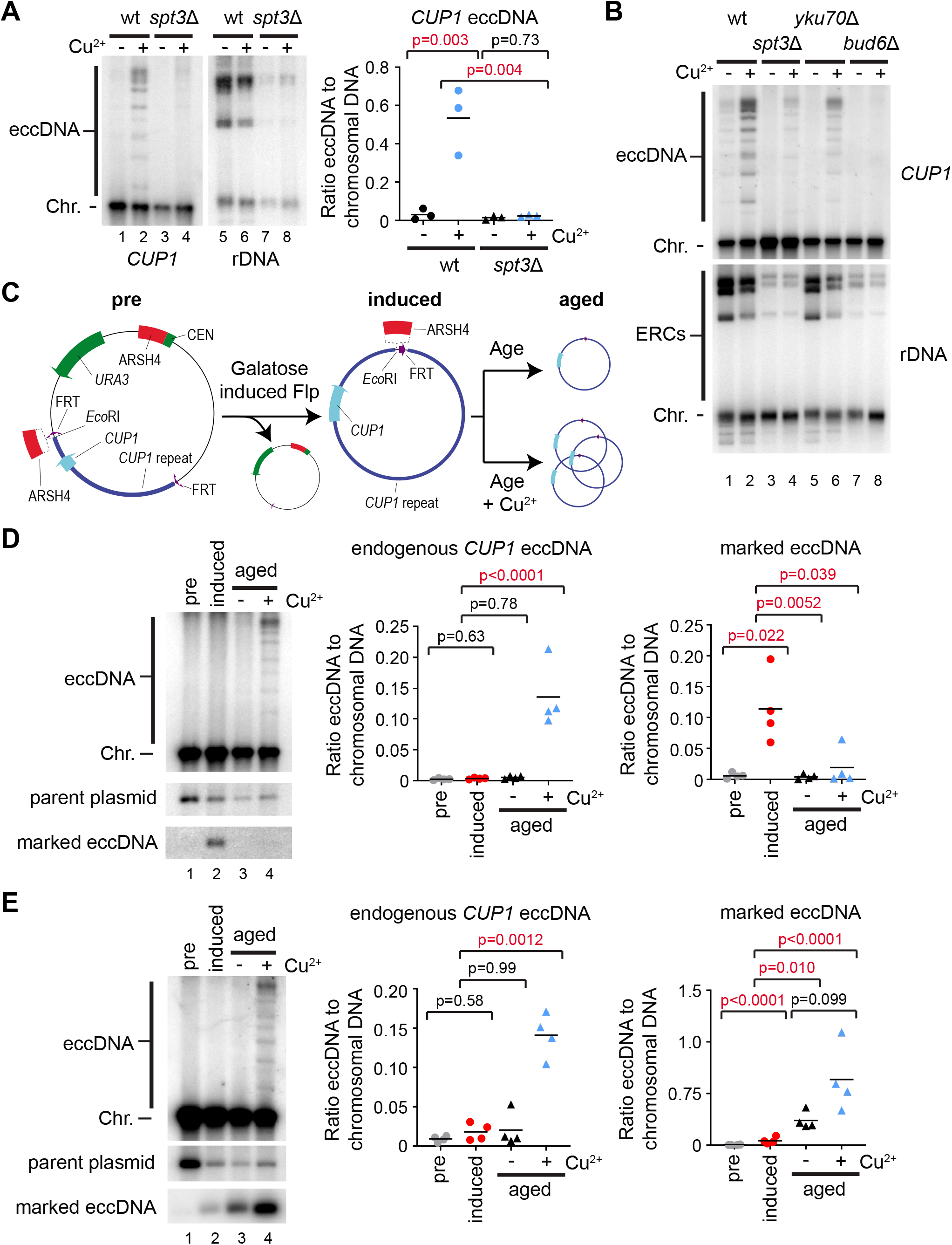
Asymmetric inheritance is necessary but not sufficient for eccDNA accumulation. A: Southern blot analysis of *CUP1* eccDNA and rDNA-derived ERCs in wild type and *spt3*Δ cells aged for 24 hours in the presence or absence of 1mM CuSO_4_, performed and analysed as in Figure 1B, n=3. B: Southern blot showing *CUP1* eccDNA and ERCs for wild type, *spt3*Δ, *yku70*Δ and *bud6*Δ cells aged for 24 hours in the presence or absence of 1mM CuSO_4_. C: Schematic representation of method to form and quantify marked *CUP1* eccDNA. Briefly, galactose-inducible Flp recombinase is used to excise a marked *CUP1* eccDNA with an *Eco*RI site absent from endogenous *CUP1* repeat region. Distribution of the marked eccDNA and endogenous species after 24 hours of ageing can be quantified by Southern blot. A variant carrying an additional ARSH4 sequence within the marked eccDNA was also constructed as shown. D: Marked eccDNA analysis: MEP wild-type cells cured of 2μ were transformed with one plasmid carrying a *CUP1* repeat and an *Eco*RI site flanked by FRT sites, and a second plasmid expressing Flp from a galactose-responsive promoter. Cells were grown overnight in selective media to maintain plasmids and containing sucrose and raffinose as carbon sources (lane 1, pre). Flp expression was induced by addition of galactose for 4 hours, then cells were labelled, inoculated in 3 cultures and grown in non-selective glucose media for 2 hours after which 1 culture was harvested (lane 2, induced). 1mM CuSO_4_ was then added to 1 culture, and both cultures were grown for 24 hours at 30°C before harvesting (lanes 3-4, aged). *Eco*RI-digested genomic DNA was analysed by Southern blot and probed for the *CUP1* repeat to reveal all species shown. Marked and endogenous *CUP1* eccDNA was quantified, band intensities were log-transformed to fulfil the assumptions of a parametric test and analysed by one-way ANOVA, n=4 biological replicates. E: Modification of marked eccDNA experiment: the ARSH4 origin of replication was cloned into the marked *CUP1* plasmid between the *Eco*RI site and the *CUP1* repeat sequence such that ARSH4 remains in the marked *CUP1* eccDNA after Flp recombination (see schematic in C). Experiment was performed exactly as in D.

We considered two mechanisms by which asymmetric segregation could be important for eccDNA accumulation. Firstly, the small population of *CUP1* eccDNA known to exist in young cells could be maintained by asymmetric segregation and amplified by replication since the *CUP1* repeat contains a replication origin [9]; this model, which is equivalent to the ERC amplification pathway, requires efficient replication that exceeds the loss rate of eccDNA by segregation to daughter cells. Alternatively, eccDNA could form frequently and be further concentrated in aged cells by asymmetric segregation; this model does not require efficient replication of eccDNA as long as the *de novo* formation rate exceeds the loss rate.

To directly test whether *CUP1* eccDNA can be maintained and amplified across ageing, we designed an inducible system to generate marked *CUP1* eccDNA in young cells (Figure 3C). A single *CUP1* repeat with an additional *Eco*RI site absent from the endogenous *CUP1* sequence was flanked by FRT sites for Flp recombinase and cloned into a stable CEN plasmid. This was transformed into a MEP strain previously cured of 2μ (a parasitic plasmid that constitutively expresses Flp), along with a second plasmid expressing Flp under a galactose-responsive promoter. Cells were pre-grown in raffinose/sucrose (Figure 3C, pre) before a 4 hour pulse with galactose to induce Flp expression and form the marked *CUP1* eccDNA (Figure 3C, induced). Labelling and recovery in glucose were then performed as normal before estradiol addition and ageing for 24 hours in glucose media with or without copper (Figure 3C, aged). After the recovery period, 11±6% of the plasmid had recombined to form the marked *CUP1* eccDNA, a low but unambiguously detectable level (Figure 3D, lane 2). After 24 hours of ageing in the presence or absence of copper the marked eccDNA was undetectable in almost all samples, despite a clearly detectable accumulation of eccDNA from the endogenous chromosomal *CUP1* locus (Figure 3D, lanes 3 & 4). This shows that *CUP1* eccDNA is not perfectly retained during ageing, and so in the absence of *de novo* formation *CUP1* eccDNA is lost.

If asymmetric segregation is sufficient for ERC accumulation, why is *CUP1* different? One possibility is that the origins of replication in each *CUP1* repeat are inefficient. We therefore constructed a variant of the marked *CUP1* eccDNA containing a high-efficiency ARSH4 (ARS209) origin (Figure 3C). In contrast to native *CUP1* eccDNA, this ARS-*CUP1* eccDNA was maintained and amplified across ageing (Figure 3E), showing that the defect in *CUP1* eccDNA maintenance stems from the replication efficiency of the eccDNA being lower than the rate of loss to daughter cells. The ARS-*CUP1* eccDNA was consistently amplified to higher levels in cells aged in the presence of copper although this effect did not reach significance; this is coherent with the requirement for SAGA component Spt3 in *CUP1* eccDNA maintenance (Figure 3A,B), as Spt3 is known to be recruited to the *CUP1* promoter on transcriptional induction, as well as to the *GAL1* promoter [58, 59]. However, this difference is small (~2-fold) relative to the differential accumulation of endogenous *CUP1* eccDNA in the presence or absence of copper (~10-fold).

Together these experiments show that *CUP1* eccDNA is subject to asymmetric segregation, but this is insufficient for maintenance as *CUP1* eccDNA is not efficiently replicated. *CUP1* eccDNA must therefore be formed at a high enough rate during ageing to offset losses through segregation to daughter cells.

### CUP1 *eccDNA is formed through DSB processing by Sae2 and Mre11*

Various mechanisms have been proposed for eccDNA formation involving repair of single or double strand breaks that are either linked to DNA replication or not (reviewed in [60]). To delimit the eccDNA formation mechanism, we analysed eccDNA accumulation in a selection of DSB processing mutants.

Many DSB processing mutants have shortened lifespans and cannot be aged sufficiently to show eccDNA accumulation [61, 62], however we characterised a small panel of informative DSB repair mutants (*dnl4*Δ, *exo1*Δ, *sae2*Δ) that aged effectively in the MEP system (Figure 4A). Firstly, loss of Dnl4, a critical DNA ligase for NHEJ, did not substantially alter *CUP1* eccDNA or ERC levels (compare lanes 4 and 12 to wild-type controls 2 and 10). Secondly, loss of exonuclease Exo1, which is involved in long-range DNA end resection as well as degradation of stalled or reversed replication forks, had no effect on *CUP1* eccDNA but substantially increased ERC levels (compare lanes 6 and 14 to wild-type controls). Thirdly, the absence of end-resection factor Sae2 caused a major reduction in *CUP1* eccDNA, but had little effect on ERCs (compare lanes 8 and 16 to wild-type controls).

**Figure 4:**
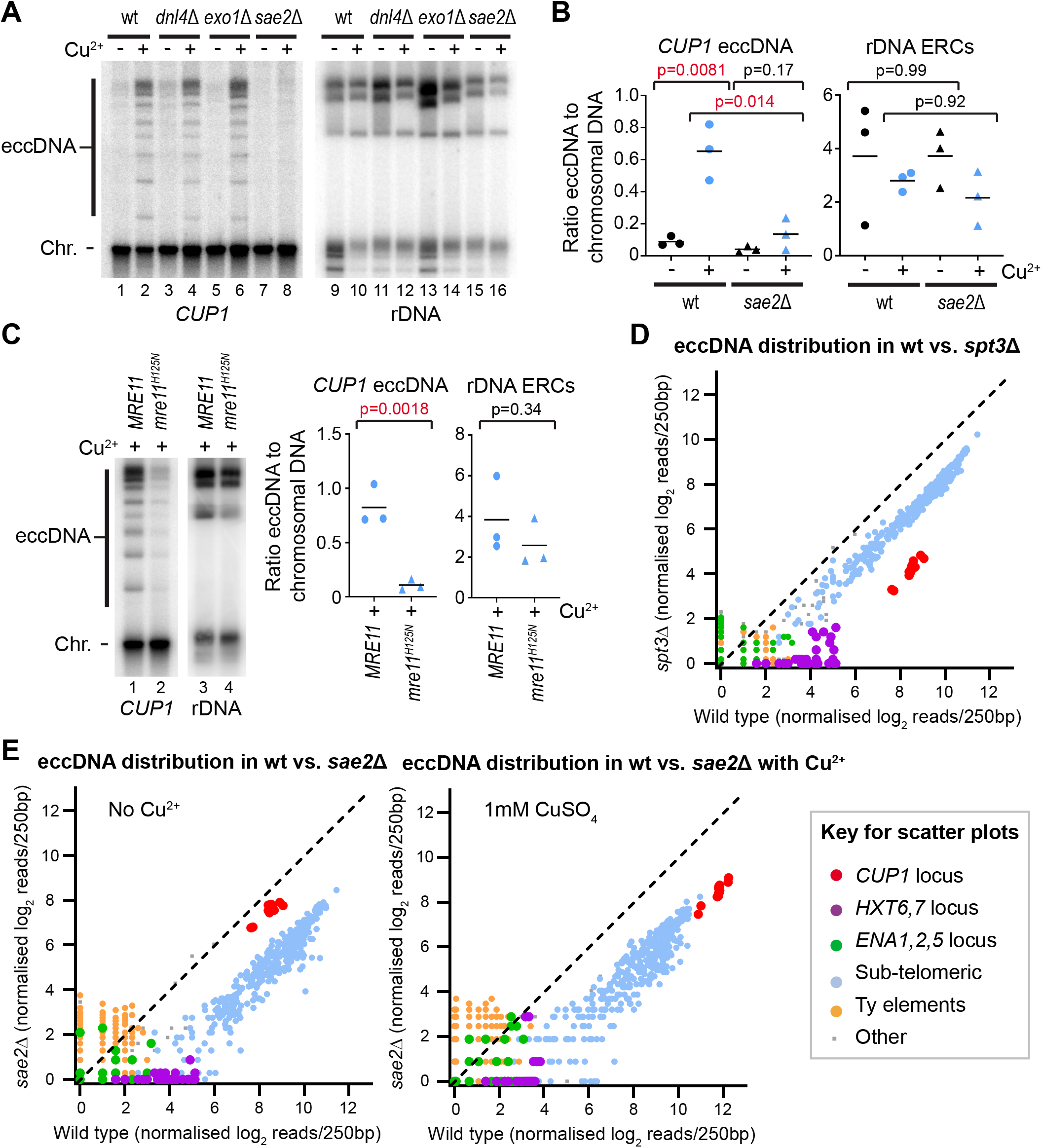
Sae2 and Mre11 nuclease activity are required for eccDNA formation. A: Southern blot analysis of *CUP1* eccDNA and rDNA-derived ERCs in wild type, *dnl4*Δ, *exo1*Δ and *sae2*Δ cells aged for 24 hours in the presence or absence of 1mM CuSO_4_, performed as in Figure 1B. B: Analysis of *CUP1* eccDNA and rDNA-derived ERCs in wild type and *sae2*Δ cells aged for 24 hours in the presence or absence of 1mM CuSO_4_, performed and analysed as in Figure 1B, n=3, quantification is derived from 3 separate clones of *sae2*Δ with 3 different selectable markers. C: Analysis of *CUP1* eccDNA and rDNA-derived ERCs in wild type and *mre11^H125N^/mre11*Δ aged for 24 hours in the presence of 1mM CuSO_4_, performed and analysed as in Figure 1B, n=3. D: REC-seq analysis comparing wild-type and *spt3*Δ cells aged for 24 hours. Samples were digested with restriction enzymes that leave the 2μ DNA intact, then read counts normalised based on 2μ counts in eccDNA and matching total DNA libraries such that eccDNA counts are relative to an equivalent quantity of total chromosomal DNA in each sample. Reads are quantified in 100bp windows spanning the genome (excluding rDNA, mitochondria, 2μ and *UBC9* locus), and coloured by feature according the key below. E: REC-seq analysis of *sae2*Δ cells compared to wild type in the absence (left) or presence (right) of 1mM CuSO_4_. Experiment and analysis as in D.

Sae2 is important for initiating DNA end resection from DSBs, particularly when DNA ends are blocked or cannot be processed (reviewed in [63]), and the importance of this factor for *CUP1* eccDNA formation suggests a formation mechanism involving DNA damage. In contrast, ERCs are known to form through a replication fork stalling mechanism that is consistent with the increased ERC levels in *exo1*Δ cells where stalled fork degradation is impaired. Therefore, the observation that *CUP1* eccDNA but not ERC accumulation is dependent on Sae2 strongly suggests that these species do not form by the same mechanism in aged cells. Usefully, the normal accumulation of ERCs in *sae2*Δ demonstrates that reduced *CUP1* eccDNA cannot simply be attributed to an ageing defect, as this would also diminish ERC levels. To ensure that this critical phenotype was reproducible, we generated two further *sae2*Δ mutants in the MEP background with different markers and consistently observed the same phenotype (Figure 4B).

Sae2 mediates end resection, stimulating the nuclease activity of Mre11 to generate a short single stranded 3’ end, and allows access to long-range resection activities of Exo1 and Dna2/Sgs1. Mre11 is a member of the Mre11-Rad50-Xrs2 (MRX) complex, and lifespan is severely compromised in *mre11*Δ cells just as for previously described *rad50*Δ mutants (data not shown) [62]. However, MRX has important functions in DSB repair beyond resection and we observed that cells carrying the nuclease deficient *mre11^H125N^* allele behaved similarly to wild type during ageing. Importantly, *mre11^H125N^* cells are defective in *CUP1* eccDNA accumulation but have little defect in ERC accumulation (Figure 4C), mirroring the *sae2*Δ phenotype (Figure 4A,B). *sgs1*Δ and particularly *sgs1*Δ *exo1*Δ mutants show variable but severe lifespan defects and therefore the importance of long-range resection could not be assessed. Nonetheless, these results show that DSB formation and resection are critical steps in *CUP1* eccDNA formation.

To reveal the locus specificity of Sae2 function in eccDNA formation, we performed REC-seq in *sae2*Δ mutants, and also in *spt3*Δ mutants to allow comparison between defects in eccDNA formation and retention. Global effects on eccDNA levels are not captured by our original REC-seq method as there is no way to normalise read counts between samples. To provide a fixed point for normalisation, we replaced one of the restriction enzymes in the REC-seq protocol with *SmaI*, which cleaves the rDNA but not the 2μ element. 2μ is a high copy circular plasmid present in all laboratory yeast strains, and can be readily detected in both eccDNA and total input DNA sequencing data. We sequenced total DNA and eccDNA from each sample and normalised eccDNA to total DNA based on 2μ read counts (the strategy is depicted in Figure S4). This modification entails a compromise as 2μ reads make up ~90% of the read count in these REC-seq libraries, dramatically reducing the signal to noise ratio from other regions, however, most eccDNA species remain detectable.

Using this modified REC-seq method, we observe that all eccDNA species are under-represented in aged *spt3*Δ cells, albeit to a variable extent, with eccDNA from the *CUP1* and *HXT6-HXT7* loci showing the greatest difference (>10 fold), while sub-telomeric circles and Ty elements are more modestly affected (~4 fold) (Figure 4D). SAGA has been previously shown to bind promoters including *CUP1* on transcriptional activation [58, 64], however the importance of SAGA for mother cell retention of *CUP1* eccDNA in the absence of copper, as well as in retaining diverse other eccDNA species, suggests that SAGA acts on eccDNA irrespective of transcription.

In contrast, the eccDNA accumulation phenotype of *sae2*Δ cells is complex. Wild-type cells aged in the presence of copper contain ~8-fold more *CUP1* eccDNA than *sae2*Δ cells (Figure 4E right panel, red spots), which is in accord with the Southern blot data (Figure 4A,B). However, this enrichment is not observed for cells aged in the absence of copper (Figure 4E left panel, red spots). This interesting difference suggests that the Sae2-dependent mechanism of *CUP1* eccDNA formation is initiated by transcription, but that a background Sae2-independent mechanism forms *CUP1* eccDNA at a basal level in un-induced cells. We also note that Ty-element eccDNA is not reduced in the absence of Sae2, consistent with the previously proposed mechanism of Ty element eccDNA formation through direct recombination between terminal repeat regions [65] (Figure 4E, orange dots). However, eccDNA from sub-telomeric circles, the *HXT6-HXT7* locus, and to some extent the *ENA1-ENA2-ENA5* locus, are also Sae2 dependent, suggesting that this mechanism of eccDNA formation acts at multiple loci (Figure 4E blue, purple and green dots respectively).

These data reveal that multiple eccDNA species form through a DSB processing pathway involving Sae2, including the transcriptionally induced *CUP1* eccDNA. However, an Sae2-independent pathway is also active, giving rise to ERCs as well as the basal level of *CUP1* eccDNA in non-copper treated cells.

### Mus81 is required for CUP1 eccDNA formation in young and aged cells

DSB resection alone does not yield eccDNA, which most likely requires intrachromosomal strand invasion at a homologous sequence (notably all the eccDNA species reproducibly detected by REC-seq are bounded by homologous regions). This recombination intermediate must be actively processed by resolvase enzymes to release circular DNA. We therefore asked which of the three known yeast resolvase enzymes Mus81-Mms4, Yen1 or Slx1-Slx4 are required. *mus81*Δ cells aged in copper had dramatically reduced *CUP1* eccDNA compared to wild type, whereas *CUP1* eccDNA levels in *yen1*Δ and *slx4*Δ mutants were indistinguishable from wild type, indicating that Mus81 is the primary resolvase for eccDNA formation (Figure 5A, lane 4).

**Figure 5:**
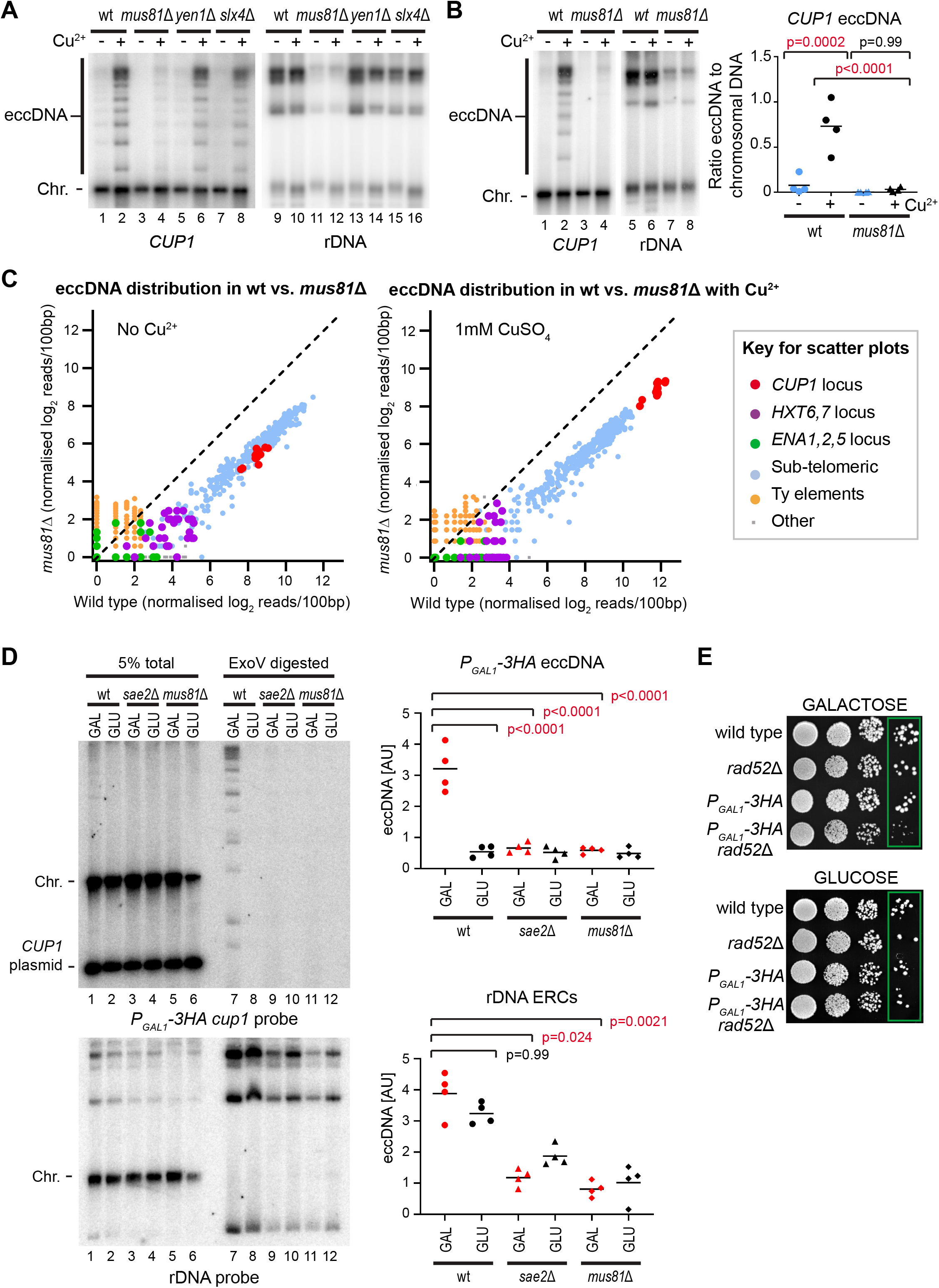
Mus81 is required for eccDNA formation in old and young cells. A: Southern blot analysis of *CUP1* eccDNA and rDNA-derived ERCs in wild type, *mus81*Δ, *yen1*Δ and *slx4*Δ cells aged for 24 hours in the presence or absence of 1mM CuSO_4_, performed as in Figure 1B. B: Analysis of *CUP1* eccDNA and rDNA-derived ERCs in wild type and *mus81*Δ cells aged for 24 hours in the presence or absence of 1mM CuSO_4_, performed and analysed as in Figure 1B, n=4. C: REC-seq analysis of *mus81*Δ cells compared to wild type in the absence (left) or presence (right) of 1mM CuSO_4_. Experiment and analysis as in Figure 4D. D: Southern blot analysis of eccDNA from the 17 copy *P_GAL1_-3HA cup1* tandem repeat in non-age selected BY4741 haploid cell background lacking MEP modifications. *P_GAL1_-3HA* wild-type, *sae2*Δ and *mus81*Δ cells were pre-grown on YP Raffinose before 6 hour induction with 2% galactose or 2% glucose. Genomic DNA was digested with *XhoI* then 95% of the sample further digested with ExoV and ExoI. 5% total DNA (lanes 1-6) and 95% ExoV digested material (lanes 7-12) are shown. These cells contain an additional pRS316-*CUP1* plasmid to complement the loss of active chromosomal *CUP1* genes, labelled as *CUP1* plasmid. Signals from same membrane stripped and re-probed for rDNA show ERC species. E: Colony formation assay performed on *P_GAL1_-3HA* wild-type and *rad52*Δ cells along with BY4741 wild type and *rad52*Δ controls. Cells were pre-grown as above on YP raffinose then serial dilutions from 10^4^ to 10^1^ cells spotted on YPD and YPGal plates, which were grown at 30° until control cells had formed equivalent sized colonies (2-3 days).

Reduced eccDNA formation in *mus81*Δ was highly reproducible (Figure 5B), however we also observed a pronounced reduction in ERC levels (Figure 5B lanes 5-8), and bud scar counting revealed that *mus81*Δ cells aged in copper are younger than wild-type cells (wild-type + Cu^2+^: 12.2±3.7, versus *mus81*Δ + Cu^2+^: 10.5±4.2), raising questions as to whether the observed *mus81*Δ phenotype actually stems from an ageing defect. Interestingly, REC-seq data comparing *mus81*Δ cells to wild type was very similar to that of *sae2*Δ cells (compare Figure 5C to Figure 4E), showing a reduced accumulation of *CUP1*, sub-telomeric circles and other species except for Ty-element eccDNA (Figure 5C), consistent with both proteins acting in the same pathway of eccDNA formation. The only notable difference is that *CUP1* eccDNA in the absence of copper is less abundant in *mus81*Δ than in wild type cells, whereas the same is not true for *sae2*Δ.

The similarity between *mus81*Δ and *sae2*Δ REC-seq profiles lead us to question the contribution of ageing to eccDNA accumulation phenotypes given that *mus81*Δ has a slow ageing phenotype. In theory, lower eccDNA levels may result not only from reduced formation, but also from cells being younger when sampled or, more subtly, from asymmetric segregation problems increasing eccDNA loss to daughter cells (as in *yku70*Δ [57]). We therefore set out to validate key aspects of the eccDNA formation mechanism in unselected populations of young cells. This would reduce age effects as the population is overwhelmingly young, and avoid segregation effects as any eccDNA formed can be detected irrespective of whether it ends up in a mother or a daughter cell. Furthermore, we used haploid cells lacking the MEP system and without biotin labelling to rule out any contribution of these experimental tools to the observed phenotype.

To enrich for eccDNA in unselected young populations, we digested genomic DNA samples with ExoV, but even after enrichment the detection of *CUP1* eccDNA was unreliable and could not be used in quantitative experiments. In contrast, detection of eccDNA from the *P_GAL1_-3HA cup1* allele (Figure 1D) proved robust over multiple experiments and provided a reliable system to test eccDNA formation. Induction of the *P_GAL1_* promoter using a 6 hour galactose pulse resulted in formation of eccDNA from the *P_GAL1_-3HA cup1* allele, showing that transcriptional activation of the locus causes eccDNA formation in young cells (Figure 5D upper panel lanes 7,8). In contrast, formation of eccDNA from this locus in cells lacking Sae2 or Mus81 was not detectable, in accord with our experiments in aged cells (Figure 5D upper panel lanes 9-12). This difference could not be attributed to a defect in transcriptional induction from the *P_GAL1_* promoter as induction was similar in all three strains (Figure S5). Curiously, we also observed that ERC levels were reduced in *sae2*Δ cells as well as in *mus81*Δ, albeit to a much lesser extent than for *P_GAL1_-3HA* eccDNA (Figure 5D lower panel). This suggests that the Sae2-dependent pathway can contribute to ERC formation; non-coding RNAs are important in rDNA recombination and it is possible that ERCs form through both transcriptionally-induced and non-transcriptionally-induced pathways that are more or less prominent in different media conditions [66]. Nonetheless, this experiment clearly demonstrates that the transcriptionally stimulated formation of eccDNA occurs through an Sae2- and Mus81-dependent DSB processing pathway, and that environmentally stimulated eccDNA formation from the *CUP1* locus is not an indirect effect of ageing in general or the experimental details of the MEP system.

Finally, our data suggests that transcriptional activation of the *CUP1* or *P_GAL1_-3HA* allele causes DSB formation, but does not exclude other options such as transcription affecting recombination outcome. To provide a more direct insight into DSB formation at this locus, we took advantage of the high efficiency of recombination in the *P_GAL1_-3HA cup1* strain. We reasoned that if DSB repair is blocked by deletion of *RAD52* in these cells, then transcriptionally-induced DSB formation should cause a prolonged arrest that would manifest in slow colony growth. We therefore plated *P_GAL1_-3HA cup1* and *P_GAL1_-3HA cup1 rad52*Δ cells on galactose and glucose plates, along with wild type and *rad52*Δ controls that should not undergo transcriptionally-induced DSB formation under these conditions. As predicted, *P_GAL1_-3HA cup1 rad52*Δ colonies grew very slowly on galactose relative to *P_GAL1_-3HA cup1 RAD52*, whereas growth was equivalent on glucose, and the *rad52*Δ mutation alone had no effect (Figure 5E, particularly the individual colonies highlighted in green boxes). This shows that DSBs are formed in response to transcription of the *P_GAL1_-3HA cup1* allele.

Overall, these data confirm that strong transcriptional induction can cause DSBs that are processed by Sae2, Mre11 and Mus81 to yield eccDNA.

## Discussion

Here, we have described the reproducible formation of eccDNA from the *CUP1* locus in response to transcriptional activation of the *CUP1* gene. *CUP1* eccDNA accumulates reproducibly to high levels through a combination of frequent formation events and retention of eccDNA in ageing cells.

### A mechanism for the formation of recurrent eccDNA

Formation of *CUP1* eccDNA has a major dependence on Sae2 and on Mre11 nuclease activity. These are processing factors involved in DSB repair by homologous recombination (recently reviewed in [67]). Sae2/Mre11 dependence therefore indicates initiation of recombination from a DSB, and argues against eccDNA formation through resolution of re-replication structures by homologous recombination, or excision of single-stranded DNA (ssDNA) loops by mismatch repair as proposed for microDNA, neither of which process is expected to be Sae2-dependent [43, 68]. Rather, our results are coherent with an intrachromosomal homologous recombination process, as proposed for formation of telomeric circles and ERCs [13, 69] (Figure 6).

**Figure 6:**
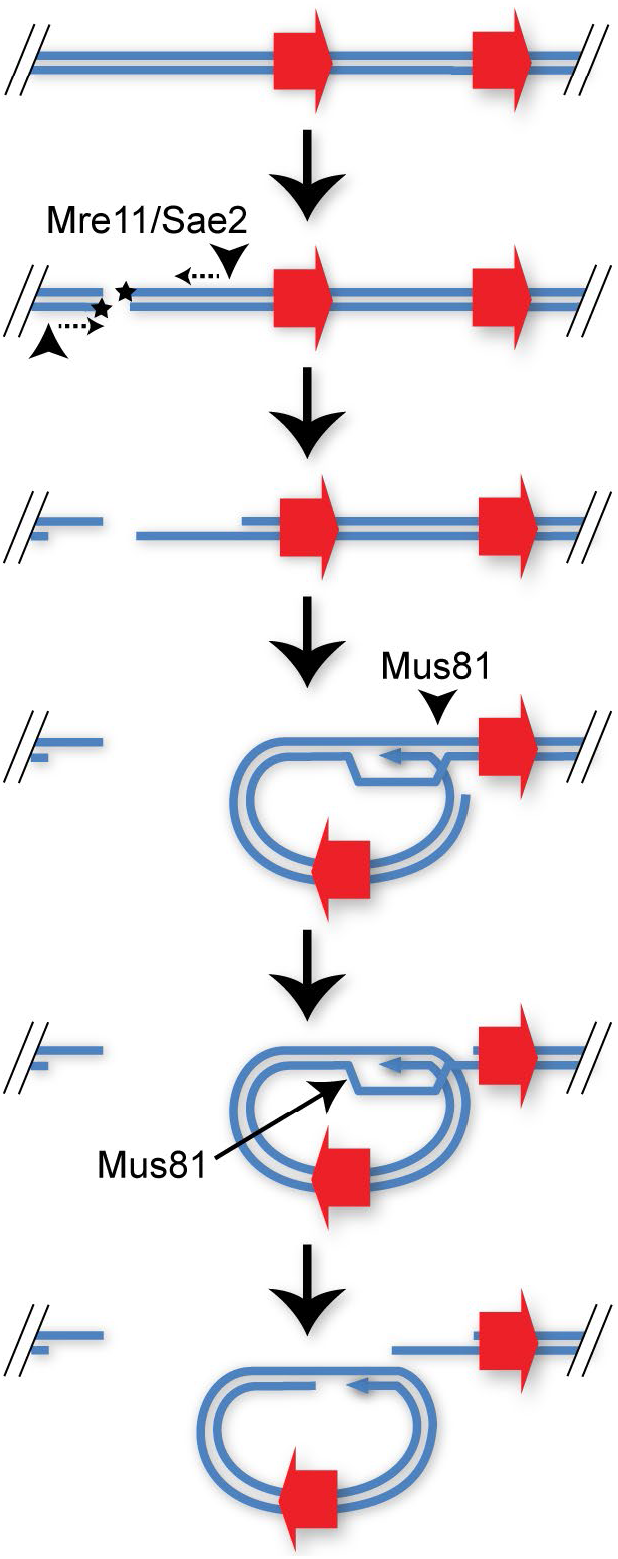
Proposed mechanism of eccDNA formation from DNA DSBs. A DSB is initially resected by Mre11 stimulated by Sae2, then invades a homologous sequence in the same chromosome. The D-loop formed is cleaved by Mus81 at two separate points, which, after ligation, yields an eccDNA and a DSB with a matching gap.

Excision of eccDNA requires two additional strand cleavages beyond the initiating DSB, and this is not a favoured outcome of intrachromosomal strand invasion (Figure 6). Helicases such as Sgs1 will dissolve D-loops formed by strand invasion to minimise rearrangements during DSB repair [70], and the activity of structure specific endonuclease enzymes capable of cleaving the nicked Holliday Junctions in a D-loop is tightly restricted to G2/M (reviewed in [71]). Nonetheless, an endonuclease activity is required and we find Mus81, which has a strong preference for cleaving nicked Holliday Junction intermediates, to be critical for eccDNA formation. This shows that eccDNA must be processed from DSBs present in late G2 or M.

The source of the DSB remains to be determined. Although transcription influences rDNA recombination, this involves a constant rate of DSB formation with a variable outcome [66], whereas the *P_GAL1_-3HA* system has a variable rate of DSB formation depending on transcription (Figure 5E). Topoisomerase II activity at highly transcribed sites can cause DSBs [72], or replication forks colliding with R-loops can be processed to DSBs, though it is not clear that Sae2 would be required for processing the latter (reviewed in [73]). R-loops may also be directly involved as Sae2 and Mre11 can remove R-loops via a DSB intermediate that could itself undergo intrachromosomal recombination [74]. Whichever the mechanism, transcriptional stimulation of any recombination process has the potential to bias recombination patterns depending on the gene expression pattern of the cell.

Previous studies have linked ERC formation to DSBs at stalled replication forks [62, 75], which appears separable from the transcription-induced Sae2-dependent *CUP1* eccDNA formation mechanism. However, there is clearly crossover between these mechanisms as *CUP1* eccDNA levels in non-copper treated cells are Sae2-independent (Figure 4E) while ERCs in log phase cells on YPRaf are partially Sae2-dependent (Figure 5D). We therefore suggest that both ERCs and *CUP1* eccDNA, and most likely other eccDNA, form both from stalled replication forks and from transcriptionally induced DSBs, with the balance between these mechanisms being variable dependent on age and environment. In contrast, Mus81 appears to be universally important, which would be consistent with Mus81 resolving the circular intermediate formed by intrachromosomal recombination initiated by either transcriptional or replication-induced DSBs.

### eccDNA formation enhances genome plasticity

The *de novo* formation of eccDNA provides various pathways for adaptation. Even in the absence of re-integration, an eccDNA with an active replication origin can amplify within a population through asymmetric segregation and selection. The presence of a high copy but heterogeneously distributed eccDNA means that all possible gene amplifcation levels are present in the population, so if survival under a particular challenge requires a certain range of copy numbers, some cells within the population should always survive. However, this enhanced adaptibility must be offset against negative effects caused by overexpression of other genes on eccDNA in the absence of the environmental challenge.

In this context it is interesting that budding yeast effectively follow two strategies. By retaining eccDNA in the ageing sub-population, the majority of young cells are able to grow unimpeded while the aged cells take on both the increased adaptability, but also the fitness cost, of maintaining eccDNAs. We and others have previously demonstrated an enhanced adaptive capacity within ageing yeast, and the accumulation of additional eccDNA likely represents an additional facet of this [76, 77]. Because eccDNA is retained in the mother it may only benefit the mother rather than any offspring. However, the machinary that retains eccDNA in the mother cell could easily be regulated under environmental stress conditions, and indeed donations of eccDNA from mother to daughter have been observed [32].

Transcriptional stimulation of eccDNA formation biases towards genes that are highly expressed in the current environment, and if inducible, this implies (though does not necessitate) that those genes have evolved to be expressed under the given environment because they are useful, and may therefore benefit from further copy number amplification. Of course not all highly expressed genes would be usefully amplified, but the yeast genome is broadly stratified into housekeeping and environmentally responsive genes based on TATA box dependence among other factors [78, 79]. It is therefore possible that a mechanism which targets DSBs to TATA-box promoters would preferentially induce eccDNA formation at highly expressed, inducible non-housekeeping genes: for example, Topoisomerase II is targeted to TATA-box promoters and can cause DSBs when the reaction is not completed [80, 81]. In this scenario, the stimulation of eccDNA formation at inducible genes that are highly expressed in the current environment makes use of information embedded in transcription patterns to direct further adaptive events, pointing to a transcription-driven mechanism conferring increased robustness/fitness through enhanced genome plasticity.

## Materials and Methods

### Strains and media

Yeast strains were constructed by standard methods and are listed in Table S1, oligonucleotide sequences are given in Table S2 and plasmids given in Table S3. Most experiments were performed in synthetic complete media (2% glucose, 0.67% yeast nitrogen base with ammonium sulphate, complete amino acid mix), to which CuSO_4_ was added from a 1 M stock solution as required. Where noted, cells were grown in YP media (2% peptone, 1% yeast extract) with sugar at 2% final volume, pre-cultures for experiments in YP were grown in YP with 2% raffinose. In all cases, cells were grown at 30°C with shaking at 200 rpm. Media components were purchased from Formedium and all media was sterilised by filtration. For all experiments, cells were brought to log phase through a 2-step culture process: cells were inoculated in 4ml media and grown for 6-8 hours then diluted in 25ml or more media and grown for 16-18 hours to 0.2-0.8×10^7^ cells/ml before further manipulation.

### MEP cell labelling and purification

For biotin labelling, 0.25×10^7^ cells per sample were harvested by centrifugation (15 s at 13,000 g), washed twice with 125 μl PBS and re-suspended in 125 μl of PBS containing ~3mg/ml Biotin-NHS (Pierce 10538723). Cells were incubated for 30 min on a wheel at room temperature, washed once with 100 μl PBS and re-suspended in 100 μl PBS, then inoculated in required media at 2×10^4^ cells/ml and allowed to recover for 2 hours at 30° before addition of 1μM β-estradiol (Sigma E2758) and CuSO_4_ as required. For ageing in YPD or YPGal, the same protocol was followed but no recovery period was included – as previously noted this prevents culture saturation without detectable contamination of young cells by 24-48 hours [31]. Cells were harvested by repeated centrifugation for 1 min, 4600 rpm in 50 ml tubes and immediately fixed by resuspension in 70% ethanol and stored at −80°C.

Percoll gradients (1-2 per sample depending on final OD) were formed by vortexing 1.16ml Percoll (Sigma P1644) with 42 μl 5 M NaCl, 98 μl water in 2 ml tubes and centrifuging 15 min at 15,000 g, 4 °C. Cells were defrosted, washed with 25 ml cold PBSE and re-suspended in a minimal amount of PBSE and layered on the pre-formed gradients. Gradients were centrifuged for 4 min at 2,000 g, then the upper phase and brown layer of cell debris removed and discarded. 1 ml PBSE was added, mixed by inversion and centrifuged 1 min at 2,000 g to pellet the cells, which were then re-suspended in 1ml PBSE (re-uniting samples split across two gradients). 25 μl Streptavidin magnetic beads were added (Miltenyi Biotech 1010007) and cells incubated for 15 min on a wheel at room temperature. Meanwhile, 1 LS column per sample (Miltenyi Biotech 1050236) was equilibrated with cold PBSE in 4 °C room. Cells were loaded on columns and allowed to flow through under gravity, washed with 1 ml cold PBSE and eluted with 1ml PBSE using plunger. Cells were re-loaded on the same columns after re-equilibration with ~500 μl PBSE, washed and re-eluted, and this process repeated for a total of three successive purifications. 50 μl cells were set aside for quality control, while 1 μl 10% Triton X-100 was added to the remainder which were then pelleted by centrifugation and frozen or processed directly for DNA extraction.

For quality control, the 50 μl cells were diluted to 300 μl final volume containing 0.3% triton X-100, 0.3 μl 1mg/ml streptavidin 594 (Life Technologies S11227), 0.6 μl 1 mg/ml WGA-488 (Life Technologies W11261) and 0.1 μg/ml DAPI in PBS. Cells were stained for 15 min – overnight at room temperature, washed once with PBS + 0.01% Triton-X100 then re-suspended in 7 μl VectaShield (Vectorlabs H-1000). Purifications routinely yielded 80-90% streptavidin positive cells with appropriate bud-scar numbers.

### DNA extraction and Southern blot analysis

For ageing samples, cell pellets were re-suspended in 50 μl 0.34 U/ml lyticase (Sigma L4025) in 1.2 M sorbitol, 50 mM EDTA, 10 mM DTT and incubated at 37 °C for 45 min. After centrifugation at 1,000 g for 5 min, cells were gently re-suspended in 80 μl of 0.3% SDS, 50 mM EDTA, 250 μg/ml Proteinase K (Roche 3115801) and incubated at 65 °C for 30 min. 32 μl 5 M KOAc was added after cooling to room temperature, samples were mixed by flicking, and then chilled on ice for 30 min. After 10 min centrifugation at 20,000 g, the supernatant was extracted into a new tube using a cut tip, 125 μl phenol:chloroform pH 8 was added and samples were mixed on a wheel for 30 min. Samples were centrifuged for 5 min at 20,000 g, the upper phase was extracted using cut tips, and precipitated with 250 μl ethanol. Pellets were washed with 70% ethanol, air-dried and left overnight at 4 °C to dissolve in 20 μl TE. After gentle mixing, 10 μl of each sample was digested with 20U *Xho*I or *Eco*RI-HF (NEB) for 3-6 hours in 20 μl 1x CutSmart buffer (NEB), 0.2 μl was quantified using PicoGreen DNA (Life Technologies), and equivalent amounts of DNA separated on 25 cm 1% 1x TBE gels overnight at 90 V (120V for gels shown in Fig. 1). Gels were washed in 0.25 N HCl for 15 min, 0.5 N NaOH for 45 min, and twice in 1.5 M NaCl, 0.5 M Tris pH 7.5 for 20 min before being transferred to 20×20cm HyBond N+ membrane in 6x SSC. Membranes were probed using random primed probes (Table S4) in 10 ml UltraHyb (Life Technologies) at 42 °C and washed with 0.1x SSC 0.1% SDS at 42 °C, or probed in Church Hyb at 65 °C and washed with 0.5x SSC 0.1% SDS at 65 °C. For probe synthesis, 25 ng template DNA in 38 μl water was denatured for 5 min at 95 °C, chilled on ice, then 10 μl 5x labelling buffer (5x NEBuffer 2, 25 μg/ml d(N)9, 165 μM dATP, dGTP, dTTP), 1 μl Klenow exo-(NEB) and varying amounts of α[^32^P]-dCTP added before incubation for 1-3 hours at 37 °C, cleaning using a miniQuickspin DNA column (Roche), and denaturing 5 min at 95 °C before adding to hybridisation buffer. During the course of this work, we learnt that controlling the amount of α[^32^P]-dCTP added to the labelling reaction was critical; too high levels resulted in high membrane background irrespective of wash conditions and probe purification. On the activity date, 0.1-0.2μl 3000Ci/mmol α[^32^P]-dCTP (Perkin Elmer) was used, an amount that was doubled every two weeks past the activity date. Images were obtained by exposure to phosphorimaging screens (GE) and developed using a FLA 7000 phosphorimager (GE).

Bands were quantified using ImageQuant v7.0 (GE) and data analysed using the GraphPad Prism v6.05. Samples were compared by one-way ANOVA with a Tukey correction for multiple comparisons; where data contravened the assumptions of a parametric test, datasets were log transformed prior to statistical testing.

For DNA samples in Figure 5D, a larger genomic DNA preparation protocol using 20×10^7^ cells per sample was employed as described [50], final genomic DNA was dissolved in 50 μl TE at 65 °C for 15 min then 4 °C overnight. 10.5 μl 10x CutSmart, 42.5 μl water and 2 μl *Xho*I were added and samples digested for 3 hours at 37 °C. 1 μl 10x CutSmart, 1 μl RecBCD [ExoV], 1 μl ExoI, 6 μl 10 mM ATP and 3.5 μl water were then added to 47.5 μl digested DNA and incubated overnight at 37 °C. Digested DNA was phenol chloroform extracted, ethanol precipitated and resuspended in 16 μl TE. These samples along with 2.5 μl *Xho*I digested total DNA, were separated and probed as above.

### Northern analysis

Northern analysis was performed as previously described [82] using probes described in [50].

### Image processing and data analysis

Gel images were quantified using ImageQuant v7.0 (GE), images for publication were processed using ImageJ v1.50i, by cropping and applying minimal contrast enhancement. Statistical analysis was performed using GraphPad Prism v7.03.

### REC-seq

All reagents from NEB except where noted otherwise. 20 μl genomic DNA isolated from aged cells as above was diluted to 155 μl final volume containing 1x CutSmart and 0.75 μl RNase T1 (Fermentas), and incubated 15 min at room temperature before splitting into 3x 49 μl and 1x 3μl aliquots (the latter was diluted to 16 μl for total DNA isolation). The 49 μl aliquots were digested with 1 μl *Eag*I-HF, *Pvu*I-HF or 0.5μl *Pvu*II-HF for 4 hours at 37 °C, 0.5 μl additional enzyme was added to the *Pvu*II digest every hour. For mutant analysis, *Pvu*II was replaced with *SmaI* and this digestion was performed at 25 °C. 6 μl 10 mM ATP, 1 μl 10x CutSmart, 1 μl Exonuclease V and 1 μl Exonuclease I was added to each digest and incubated over night at 37 °C before extraction with phenol chloroform and ethanol precipitation in the presence of 1 μl GlycoBlue (Life Technologies). DNA pellets were dissolved in 45 μl 0.1x TE, then 6 μl 10x CutSmart, 6 μl 10 mM ATP, 1 μl RecBCD [ExoV], 1 μl Exonuclease I and 1 μl same restriction enzyme as previous day were added and samples again incubated over night at 37 °C before extraction with phenol chloroform and ethanol precipitation. For *Sma*I digestions, the enzyme and CutSmart were added first and digestion performed for 4 hours at 25 °C before addition of other components and shift to 37 °C for overnight incubation. Pellets were dissolved in 33 μl 0.1x TE and the 3 aliquots combined, precipitated again with ethanol and pellets dissolved in 16 μl 0.1x TE.

Exonuclease treated and total DNA samples were incubated at 37 °C for 45 min with 2 μl NEBNext DNA fragmentase and 2 μl fragmentase buffer, then 5 μl 0.5 M EDTA was added followed by 25 μl water and samples were purified using 50 μl AMPure beads and eluted with 25.5 μl 0.1x TE. 3.5 μl NEBNext Ultra II end prep buffer and 1.5 μl NEBNext Ultra II end prep enzyme were mixed in and samples incubated 30 min at 20 °C then 30 min at 65 °C. After addition of 1.25 μl 1:25 diluted NEBNext adaptor, 0.5 μl ligation enhancer and 15 μl NEBNext Ultra II ligation mix samples were incubated 15 min at 20 °C, then 1.5 μl USER enzyme was added and incubated 15 min at 37 °C. Samples were cleaned with 44 μl AMPure beads, eluted with 30 μl 0.1x TE, then cleaned again with 27 μl AMPure beads and eluted with 22.5 μl 0.1x TE.

1.25 μl library was amplified with 0.4 μl each NEBNext index and universal primers and 5μl NEBNext Ultra II PCR mix in 10 μl total volume using recommended cycling conditions with 8 cycles for total samples, 16 cycles for 48 hour aged samples and 18 cycles for 24 hour samples. These test amplifications were cleaned with 9 μl AMPure beads, eluted with 1.5 μl 0.1x TE and 1 μl run on a Bioanalyzer to determine the optimal cycle number for the final amplification. Library yield was aimed to be 2 pM, the test amplification should be equivalent to a 1:4 dilution of the final library. The remaining library (21 μl) was then amplified in a 50 μl reaction containing 2 μl each NEBNext index and universal primers and 25 μl NEBNext Ultra II PCR mix using the calculated cycle number, cleaned with 45 μl AMPure beads, eluted with 25 μl 0.1x TE, then cleaned again with 22.5 μl AMPure beads and eluted with 10.5 μl 0.1x TE.

### Sequencing and bioinformatics

Libraries were sequenced on a NextSeq500 (Illumina) in Paired End 75 bp Mid-output mode by the Babraham Institute Sequencing Facility, and basic data processing performed by the Babraham Institute Bioinformatics Facility. After adapter and quality trimming using Trim Galore (v0.5.0), REC-seq data was mapped to yeast genome R64-2-1 with 2μ sequence J01347.1 included as an additional chromosome using Bowtie2 (v2.3.2; with more stringent parameters: --no-mixed --score-min L,0,-0.2 --no-unal -X 2000). To avoid apparent enrichment of repetitive regions (https://sequencing.qcfail.com/articles/deduplicating-ambiguously-mapped-data-makes-it-look-like-repeats-are-enriched/) the resulting BAM files were subjected to de-duplication based on exact sequence identity of the first 50 bp of both Read 1 and Read 2 using a custom script (available on request). These de-duplicated by sequence files were then imported into SeqMonk v1.44.0 for analysis (https://www.bioinformatics.babraham.ac.uk/projects/seqmonk/), where they were additionally de-duplicated based on alignment position (chromosome, start and end). Reads were quantified in 20 bp bins and normalised based on the total read count in each library. For 3x*CUP1*/Δ libraries, counts were performed in 100 bp bins and data from replicate sets compared using the edgeR statistical filter implemented in SeqMonk [83].

All raw REC-seq data has been deposited at GEO under accession number GSE135542.

## Acknowledgements

We thank Kristina Tabbada and Clare Murnane in the BI Next Generation Sequencing Facility for data generation, Anne Segonds-Pichon and Simon Andrews of the BI Bioinformatics facility for statistical and bioinformatics advice, Birgitte Regenberg and Henrik Moller for helpful discussions and Tanya Paull and Dan Gottschling for reagents. Funding for JH, RH, MK and GP was from the Wellcome Trust [088335,110216], wellcome.ac.uk, for JH and FK from the Biotechnology and Biological Sciences Research Council BBSRC [BI Epigenetics ISP: BBS/E/B/000C0423], https://bbsrc.ukri.org/, and for XV from the European Commision [Erasmus+ programme], www.erasmusplus.org.uk. The funders had no role in study design, data collection and analysis, decision to publish, or preparation of the manuscript.

